# Continuous administration of the p38α inhibitor neflamapimod during the subacute phase after transient ischemia-induced stroke in the rat promotes dose-dependent functional recovery accompanied by increase in brain BDNF protein level

**DOI:** 10.1101/2020.04.29.068015

**Authors:** John Alam, Michael Krakovsky, Ursula Germann, Aharon Levy

**Affiliations:** EIP Pharma, Inc., Boston, Massachusetts, United States of America; Pharmaseed Ltd., Ness-Ziona, Israel

## Abstract

There is unmet need for effective stroke therapies. Numerous neuroprotection attempts for acute cerebral ischemia have failed and there is growing interest in developing therapies that widen the treatment initiation window and promote functional recovery through increasing synaptic plasticity. The p38α mitogen-activated protein kinase is an already proven target for acute experimental stroke intervention and was hypothesized to also contribute to neuroinflammation-mediated impairment of recovery during the subacute phase. Neflamapimod, an orally bioavailable, brain-penetrant, potent and selective small molecule p38α inhibitor was evaluated as a subacute phase stroke treatment to promote recovery in this research study. Neflamapimod administration at two clinically relevant dose levels was initiated outside of the previously characterized neuroprotection window of less than 24 hours after stroke for p38α inhibitors to rats after transient middle cerebral artery occlusion. Continuous administration of neflamapimod, starting at 48 hours after reperfusion, significantly improved behavioral outcomes assessed by the modified neurological severity score at four- and six-weeks post stroke in a dose-dependent manner. Neflamapimod also demonstrated beneficial effects on additional measures of sensory and motor function and resulted in a dose-related increase in the terminal brain-derived neurotrophic factor protein level in both the injured and uninjured brain hemisphere. Variable interleukin-1β levels were detected in the injured brain hemisphere at study termination in a subset of the animals within every test group, implying ongoing, chronic inflammation, however, no clear neflamapimod effect on interleukin-1β production was observable. The dose-related *in vivo* efficacy of neflamapimod offers the possibility of both expanding the window for initiation of therapy after stroke and for improving recovery after a completed stroke. Since neflamapimod is already in mid-stage clinical trials for Alzheimer’s disease and related dementias, the current results make it especially attractive for evaluation in a proof-of-concept clinical trial as therapeutic to promote recovery after ischemic stroke.

## Introduction

Stroke is a frequent cause of death as well as a leading cause of acquired disability worldwide and is associated with a substantial economic burden due to high costs for treatment and post-stroke care [1, 2]. Approximately 80% of strokes are ischemic in nature due to thromboembolic occlusion of a major artery or its branches, leading to a cascade of events that causes irreversible tissue injury [3]. The only approved medical treatments for acute ischemic stroke are intravenous thrombolysis with recombinant tissue plasminogen activator (TPA), resulting in recanalization of occluded vessels if applied within a short time period (up to 4.5 hours) after stroke [4], or thrombectomy, ideally performed within 6 hours after stroke symptom onset (although under certain conditions beneficial if applied within 24 hours after stroke symptom onset), involving mechanical removal of the thromboembolic material occluding the vessel lumen [5].

Based on pathological characteristics and their timing, a stroke is classified into three clinical phases, including the acute (i.e. first 48 hours after stroke onset), the subacute (from 48 hours to >6 weeks post stroke) and the chronic phase (starts at 3-6 months post stroke) [6]. The acute phase represents an opportunity to salvage threatened tissue and reduce the extent of injury, for example via reperfusion or neuroprotection [7, 8]. The subacute phase represents the recovery stage, which is characterized by brain repair initiation, so therapeutic strategies include enhancing the underlying spontaneous recovery processes, modifying inflammation, lifting diaschisis, or reducing late neuronal death [7, 8].

Numerous attempts at providing neuroprotection during the acute phase of stroke have failed and currently, there is no United States Food and Drug Administration (FDA)-approved agent for neuroprotection after ischemic stroke [9-11]. Clearly there is an urgent need for alternative, more widely applicable treatment options for ischemic stroke including therapeutics that enable expansion of the time window from stroke to treatment. While it remains important to continue to advance novel acute stroke interventions, there is also high interest in the development of novel therapies that are directed at promoting functional recovery from stroke via increasing neuronal and synaptic plasticity during the subacute phase [8, 9, 12]. The goal is to identify disease-modifying treatments that can be administered after the acute phase of stroke is complete [8, 9, 12]. It is envisioned that such neurorestorative agents are administered during the subacute and/or the chronic phase post stroke only and that they may target compromised cerebral tissue and/or surrounding intact tissue to promote brain plasticity [8, 12].

The p38 mitogen-activated protein kinase (MAPK) pathway plays a crucial role in several central nervous system (CNS) disorders, including Alzheimer’s disease, Down’s syndrome, Parkinson’s disease, spinal cord injury (SCI), and cerebral ischemia [13, 14]. Among the four p38 MAPK family members, p38α was discovered first as a stress-activated protein kinase that plays a central role in inflammation [15, 16]. It is also the best characterized isoform as a target for CNS drug discovery [17, 18]. A large variety of biological roles have been attributed to p38α in brain pathology which depend on the type and stage of the CNS disorder, the brain region, the cell type, and include modulation of pro-inflammatory cytokine (e.g. IL-1β and tumor necrosis factor α (TNFα)) production as well as signaling (e.g. in glia, microglia, astrocytes, neurons) in the brain, and orchestration of neurotoxicity, neuroinflammation and/or synaptic dysfunction [14, 17-19].

In the adult mouse, p38α is highly expressed in different brain areas including cerebral cortex, hippocampus, cerebellum, and a few nuclei of the brainstem [20]. At the subcellular level, p38α is distributed in dendrites and in cytoplasmic and nuclear regions of the cell body of neurons [20]. Activation of p38 MAPK, including p38α, signaling after experimental ischemic stroke in rodents has been demonstrated in neurons, astrocytes and microglia [21-25], and p38α has been established as a driver of neuroinflammation-mediated cell death in the acute phase of ischemic stroke [26, 27]. Therefore, several inhibitors of p38 MAPK that exhibit different potency and kinase selectivity, all of them most potently blocking p38α versus other p38 isoforms (p38β, p38γ, p38δ) have been administered during the acute phase with treatment initiation within the first 24 hours after stroke in experimental models of cerebral ischemia, and all of them have provided robust neuroprotection [25, 28-32].

While p38α has already been proven as a therapeutic target for acute stroke intervention, it is unclear whether p38α also plays a critical role in impairing functional recovery during the subacute phase of stroke which embodies brain repair initiation [6-8]. Time course analyses of phospho-p38 (i.e. the activated from of p38) expression in experimental rat stroke models reported increased phospo-p38, including phospho-p38α, immediately after cerebral ischemia within neurons and other cell types, and, despite some fluctuations observed in some of the studies, also demonstrated that phospho-p38α remained elevated and associated with neurons, as well as other cell types, for >4 days, and in one study up to 14 days, implying that p38α-activation during the subacute phase could be partly responsible for poor recovery from stroke [22, 24, 25, 31]. A one week study of a novel p38α inhibitor, the tetra-substituted thiophene VCP979, evaluated as a treatment of phothothrombotic ischemic stroke in streptozocin-induced Type 2 diabetes mellitus (T2DM) mice demonstrated beneficial efficacy resulting in neuroprotection as well as axonal/white matter remodeling within the motor cortex [32]. Whereas this study offered novel mechanistic insights for a p38α inhibitor that are relevant for brain repair, it did not address widening the window to treatment initiation since VCP979 administration started at 24 hours post stroke [32], at a time point that is within the acute phase of photothrombotic stroke in mice [33, 34].

Brain-derived neurotrophic factor (BDNF) promotes neural plasticity and recovery after stroke [35] and the proinflammatory cytokine interleukin-1 beta (IL-1β) is upregulated after ischemic stroke [36-40]. In subacute/chronic inflammatory conditions, IL-1β is known to be a key component of the inflammatory response in the brain that mediates neurodegenerative effects of inflammation on cognition and synaptic plasticity [41]. Chronic elevation of IL-1β, such as IL-1β elevation in the aging brain, suppresses BDNF production [42], and *in vitro* it has been shown that increased IL-1β inhibits BDNF effects on neuronal/synaptic plasticity via induction of p38 MAPK [43]. All the above-described research findings taken together stimulated the idea to investigate whether neflamapimod, a blood brain barrier-penetrant small molecule p38α inhibitor, may have beneficial efficacy as a subacute phase stroke treatment via promoting functional recovery through increasing synaptic plasticity.

The pyrimido pyridazine neflamapimod (International Non-proprietary Name for molecule previously code-named VX-745) is a potent, highly selective, ATP-competitive inhibitor of p38α that yields higher CNS than peripheral blood exposure after oral administration [44, 45]. It exhibits efficacy and safety profiles in preclinical species to merit ongoing clinical investigation for CNS disorder indications [45]. Neflamapimod is currently being evaluated in the clinic for its potential to reverse synaptic dysfunction in three different CNS disorders (Alzheimer’s disease, Huntington’s disease, dementia with Lewy bodies). The first two exploratory clinical studies of neflamapimod in 9 and 16 subjects, respectively, with mild cognitive impairment due to Alzheimer’s disease demonstrated that neflamapimod penetrates the blood-brain barrier in humans and exhibits disease-relevant pharmacological activity in the brain [45, 46].

In a previous research study [47], neflamapimod was evaluated in 20- to 24-month old Fischer rats with cognitive deficits resulting from IL-1β-induced impairment of synaptic plasticity [48]. Results showed that a very low dose of neflamapimod (1.5 mg/kg administered twice daily) reversed spatial learning deficits in the Morris-Water-Maze test with no effects on hippocampal IL- 1β levels, and that a 3-fold higher dose (4.5 mg/kg administered twice daily) reduced IL-1β levels in the hippocampus but did not result in pro-cognitive effects [47]. These results suggested that the neflamapimod pro-cognitive effects were mediated through decreasing IL-1β signaling in hippocampal neuron target cells, rather than through reducing IL-1β production [47]. Thus, the present study was also undertaken to extend the aged rat behavioral results by investigating these two clinically relevant neflamapimod dose levels in a subacute phase treatment experimental paradigm for their efficacy in promoting neurologic plasticity and functional recovery in experimental stroke.

In recognition of the unmet need for stroke therapies that enable a wider treatment window and show potential for promoting functional recovery via beneficial effects on synaptic plasticity, the objectives of the present study were (1) to evaluate neflamapimod at clinically relevant dose levels for its *in vivo* efficacy to promote neurologic recovery as assessed by the modified neurological severity score (mNSS) and additional behavioral tests in a rat transient middle cerebral artery occlusion (tMCAO) model; (2) to initiate administration of neflamapimod outside the known neuroprotection window for p38α inhibitors and address the possibility for widening the window for treatment post stroke induction; (3) to assess the neurogenic factor BDNF in the brain as a potential biomarker for monitoring neflamapimod effects on synaptic plasticity; and (4) to measure IL-1β levels as an inflammatory biomarker in the brain after the treatment period to gauge whether chronic inflammation may play a role in this experimental stroke model and whether a neflamapimod anti-inflammatory effect may be detectable.

## Materials and methods

### Animals and general health monitoring

Seventy-six young (3-month old) male Sprague Dawley rats (Harlan Laboratories, Israel) weighing 328 g ± 20% were included in the study. The protocol for the study was approved by the Israeli Animal Care and Use Committee (approval number IL-15-01-15) and was conducted in accordance with the Israeli guidelines that conform to the United States Public Health Service’s Policy on Humane Care and Use of Laboratory Animals. Animal handling was performed according to guidelines of the National Institute of Health (NIH) and the Association for Assessment and Accreditation of Laboratory Animal Care (AAALAC). To assess the health status of the animals throughout the study, the general health status was monitored daily and body weight was determined weekly.

### Transient middle cerebral artery occlusion

Transient middle cerebral artery occlusion (tMCAO) in the right brain hemisphere was performed on Day 1 according to the method of Longa *et al*. [49], in which a 4 cm length of 4-0 monofilament nylon suture is inserted under anesthesia through the proximal external carotid artery into the internal carotid artery and thence into the circle of Willis, effectively occluding the middle cerebral artery. Anesthesia was induced with 4% isoflurane in a mixture of 70% N2O and 30% O2 and maintained with 1.5-2% isoflurane. The surgical wound was subsequently closed, and the animals were returned to their cages to recover from anesthesia. Two hours after occlusion rats were re-anesthetized, monofilament was withdrawn to allow reperfusion, surgical wound was closed, and rats were returned to their cages.

### Neurologic scoring and behavioral evaluations

The individual performing neurologic scoring and behavioral assessments was unaware of the group assignments, thus, performed blinded evaluations. Neurologic scoring by mNSS evaluation according to Chen *et al*. [50] was performed at least one day before the Day 1 tMCAO, and on Day 2, at Week 4, and at Week 6 following tMCAO. The mNSS values included the results of a set of clinical-neurological tests, a composite of motor, sensory, reflex and balance tests and well-defined score values (summarized in Table 1 [50]), that were used to assess the effect of neflamapimod compared to vehicle control. Taking into account the different tests and score values described in Table 1, the mNSS was graded on a composite scale of 0 to 18, in which the normal, healthy animal value was 0 and the maximal deficit value after tMCAO was represented by 18 [50]. An overall mNSS value of ≥10 was predefined as an inclusion criterion for enrolling a rat with stroke into the neflamapimod treatment study.

**Table 1.**
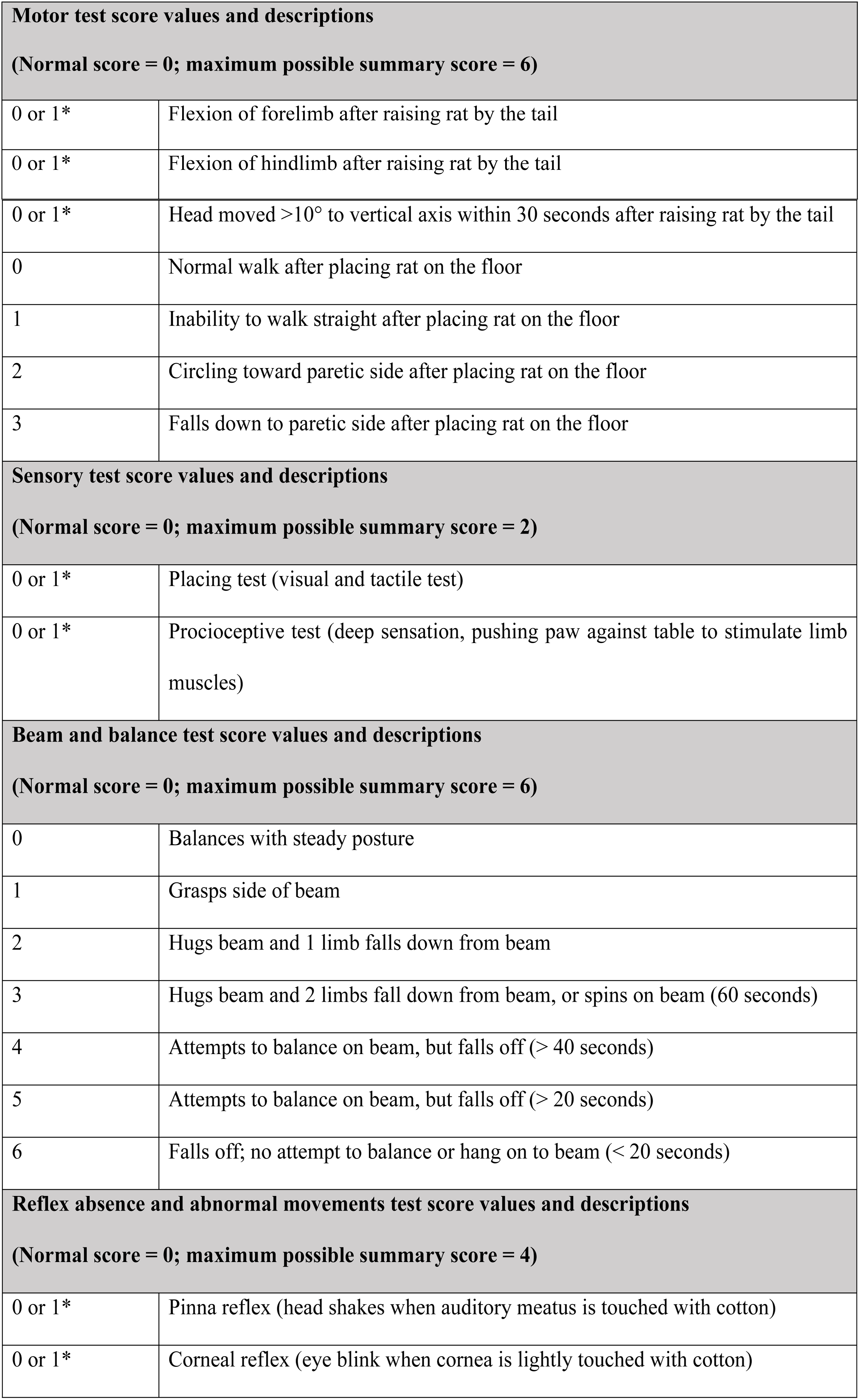
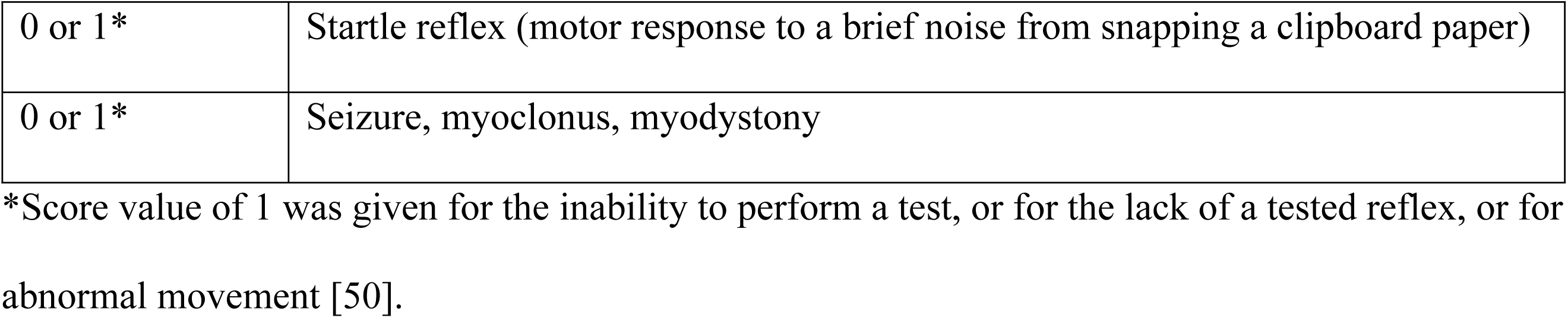
Modified Neurological Severity Score (mNSS) tests and scoring values [50].

Additional behavioral evaluations included stepping test, body swing test, and forelimb placement test that were performed before the tMCAO procedure, and at Week 4 and Week 6 following tMCAO.

A stepping test was utilized to assess forelimb akinesia according to Pharmaseed’s internal protocol. Each rat was held with its hind limbs fixed in one hand and the forelimb, not to be monitored, in the other, while the unrestrained forepaw touched the table. The number of adjusting steps of the forelimb to be monitored was counted while the animal was moved sideways along the table surface in the forehand and backhand direction over a distance of 85 cm during approximately five seconds. The stepping test was completed for both the left and right forelimbs at all time points indicated above.

For the body swing test, each rat was held approximately one inch from the base of its tail. It was then elevated to an inch above a surface of the table. The rat was held in the vertical axis, defined as no more than 10° to either the left or the right side. A swing was recorded whenever the rat moved its head out of the vertical axis to either side. Before measuring another swing, the rat must have returned to the vertical position. Twenty (20) total swings were counted. A normal rat typically had an equal number of swings to either side. Adjusted body swing scores were calculated as the difference between leftward and rightward swings (i.e. the number of rightward swings subtracted from leftward swings).

For the forelimb-placing test, the examiner held the rat close to a tabletop and scored the rat’s ability to place the forelimb on the tabletop in response to whisker, visual, tactile, or proprioceptive stimulation. Separate sub-scores were obtained for each mode of sensory input and added to give total scores (0=normal, 12=maximally impaired). Scores were given in half-point increments: whisker placing (0-2); visual placing (forward (0-2) and sideways (0-2)); tactile placing (dorsal (0-2) and lateral (0-2)); proprioceptive placing (0-2).

### Administration of test articles

Test articles, neflamapimod at two different dose levels in 1% (w/v) Pluronic F108 vehicle, or vehicle (1% (w/v) Pluronic F108) were supplied by EIP Pharma, Inc. and administered at a dosing volume of 5 ml/kg by oral gavage twice daily (7 AM and 7 PM) on 5.5 days (i.e. Sunday through Friday at 7 AM and Sunday through Thursday at 7 PM) every week for six weeks (i.e. from Day 3 until Day 42), starting at 48 hours post reperfusion.

### Sample collection and analysis

On Day 44, two days following the final dosing of test article or vehicle, rats were sacrificed by CO_2_ inhalation. Brains were harvested from all animals and samples were divided into left and right hemisphere. Samples were weighed, immediately frozen in liquid nitrogen, and stored at -80°C. For the IL-1β and BDNF analyses, tissue samples were defrosted and homogenized in 1 ml/200 mg tissue of 20 mM Tris-HCL pH 7.4 containing a protease inhibitor cocktail (50 μl/ml). Samples were centrifuged at 10,000 x g for 15 minutes at 4°C. Clear supernatants were aliquoted (150 μl/aliquot) and stored at -80°C until enzyme linked immunosorbent assay (ELISA) for IL-1β or BDNF was performed. For IL-1β analysis, ELISA was performed using the Rat IL-1 beta/IL-1F2 Quantikine ELISA Kit (R&D Systems, Minneapolis, MN) according to manufacturer’s instructions. For BDNF analysis, ELISA was performed using the Solid Phase Sandwich Quantikine ELISA Kit (96-well strip plates, R&D Systems, Canada) according to manufacturer’s instructions. For IL-1β and BDNF, standards and samples were tested in duplicates. For IL-1β assay, all the results below the lower limit of quantification (LLOQ) were assigned the LLOQ value of 5 pg/ml. For BDNF assay, all the results were above the LLOQ.

### Statistical analysis

For body weight monitoring, mNSS or other behavioral evaluations, and IL-1β levels, statistical analysis was performed by two-way analysis of variance (ANOVA) for repeated measures, followed by Bonferroni post-hoc test to adjust for the multiple comparisons. For BDNF levels, Jonckheere-Terpstra test was utilized to assess for treatment-dependent dose effect across placebo, 1.5 mg/kg, and 4.5 mg/kg dose groups. In addition, one-way ANOVA followed by Bonferroni’s multiple comparison test was performed to compare individual neflamapimod dose groups to the placebo group. One-way ANOVA was also used to compare the mNSS values at two different time points (e.g. Week 4 versus Day 2, or Week 6 versus Week 4) within an individual test group.

## Results

### Robust experimental stroke induction in rats caused significant increase in mNSS value and low mortality during the acute phase (i.e. 0-48 hours) after stroke

A frequently used method, tMCAO, was used to generate a robust experimental stroke as assessed by mNSS functional testing or mortality. Ischemia was induced on Day 1 in 76 young adult, 3-month old male Sprague-Dawley rats by occlusion of the right middle cerebral artery, which was released after 2 hours [49]. Nine animals were lost during the first day and four additional animals were lost during the second day after the procedure, resulting in ∼17% mortality during the first 48 hours post tMCAO attributable to the severity of these rat’s stroke.

On Day 2, at 24 hours after reperfusion, 67 surviving animals were subjected to neurological evaluation using the mNSS value (see Table 1 [50]) that rates neurological functioning on a scale from 0 (healthy) to 18 (maximum impairment) [51]. The measured mNSS value in individual rats rose from 0 before the Day 1 tMCAO procedure to a mean±standard deviation (SD) value of 14.0±1.4 on Day 2, with individual animal mNSS values ranging from 12 to 17, indicating successful induction of a robust experimental stroke.

On Day 3, at 48 hours post stroke, a total of 63 animals remained available for inclusion in the study and the animals (n≥20 per group at randomization) were randomized based on Day 2 mNSS values and assigned to one of three treatment groups: twice daily 1.5 mg/kg neflamapimod (i.e. 22 rats), twice daily 4.5 mg/kg neflamapimod (i.e. 21 rats), or twice daily vehicle (i.e. 20 control rats) during 5.5 days every week for a total of six weeks.

### Observations of animal health preservation including body weight gain after initiation of neflamapimod or vehicle treatment during the subacute phase after stroke

The first administration of neflamapimod treatment or vehicle control administration occurred on Day 3 at 48 hours post reperfusion. Overall animal health throughout the study was assessed by weekly body weight determinations, daily health assessment and survival observations. Three additional rats, including one animal from the 1.5 mg/kg neflamapimod dose group and two vehicle control rats, did not survive the first two weeks after initiation of dosing. Since these additional deaths were inversely related to the dose of neflamapimod, they were attributed to the severity of their stroke. These late mortalities indicate that the recovery process after a late treatment initiation in this rat tMCAO model was very slow during the first two weeks post-stroke. The 60 remaining rats (n=21 in the 1.5 mg/kg and n=21 in the 4.5 mg/kg neflamapimod groups, respectively, and n=18 in the vehicle group) completed the planned six weeks of dosing and were included in the neurologic and behavioral evaluations at Week 4 and Week 6.

Although the three experimental treatment groups were randomized based on Day 2 mNSS values, a retrospective analysis of the Day 1 mean±SD body weight data for the animals assigned to each group revealed that their body weights were also well balanced (328.5±11.3 g for the 1.5 mg/kg and 325.3±11.1 g for the 4.5 mg/kg neflamapimod treatment group, respectively, and 328.2±9.6 g for the vehicle control group) as shown in Table 2. Weekly body weight monitoring and two-way ANOVA statistics followed by Bonferroni post-hoc comparisons revealed no statistically significant differences in body weight for the three study groups at any time throughout the study. Additionally, no differences in general clinical signs were observed upon daily animal health monitoring (data not shown). The mean±SD body weight for the 3-month old rats was 327.0±10.9 g on Day 1 and all surviving animals had a mean±SD body weight 418.3±27.6 g at Week 6, and a similar body weight gain of ∼27% was observed in all three test groups throughout the study period (Table 2). These observations together with favorable results for all the daily health assessments point out that the neflamapimod and vehicle treatments were generally well-tolerated, providing no treatment-related adverse clinical signs. Moreover, these findings exclude the possibility that a difference in the general health of the rats contributed to neflamapimod treatment-mediated effects on functional recovery when compared to the effects observed in the vehicle-treated group.

**Table 2.** Body weight monitoring of the three treatment groups, including vehicle control, 1.5 mg/kg neflamapimod (NFMD), 4.5 mg/kg NFMD throughout the study.

| **Body weight**<br>**monitoring time** | **Body weight**<br>**Vehicle**<br>**Mean±SD (g)** | **Body weight**<br>**1.5 mg/kg NFMD**<br>**Mean±SD (g)** | **Body weight**<br>**4.5 mg/kg NFMD**<br>**Mean±SD (g)** |
| --- | --- | --- | --- |
| **Day 1** | 328.2±9.6 | 328.5±11.3 | 325.3±11.1 |
| **Week 1** | 309.2±36.3 | 325.7±26.8 | 321.1±27.2 |
| **Week 2** | 342.2±34.2 | 353.9±24.9 | 353.3±23.8 |
| **Week 3** | 369.2±30.7 | 378.5±23.1 | 374.8±21.4 |
| **Week 4** | 388.6±30.5 | 397.3±25.1 | 393.9±25.5 |
| **Week 5** | 403.6±31.4 | 412.4±25.6 | 407.5±21.1 |
| **Week 6** | 415.0±34.5 | 421.5±26.8 | 417.8±23.0 |

### Dose-related improvement in neurologic and behavioral mNSS after initiation of neflamapimod versus vehicle treatment during the subacute phase post stroke

Neurologic and behavioral improvements are reliable parameters for measuring the therapeutic efficacy in experimental stroke and the mNSS is one of the most commonly used neurological scales in experimental stroke studies that includes a composite of motor (muscle status and abnormal movement), sensory (visual, tactile, and proprioceptive), balance and reflex tests (see Table 1) on a scale up to 18 for rats [50, 51]. On Day 2, the mean±SD values for mNSS were similar across the groups of stroked rats, indicating that the three randomized experimental groups were balanced a day prior to treatment initiation (Fig 1A). The measured mNSS mean±SD values on Day 2 were 13.9±1.1 in the 1.5 mg/kg and 13.9±1.4 in the 4.5 mg/kg neflamapimod dose group, and 14.3±1.7 in the vehicle control group. In light of the maximum possible mNSS value of 18 (Table 1), all animals exhibited severe neurologic deficits prior to neflamapimod treatment initiation.

**Fig 1.**
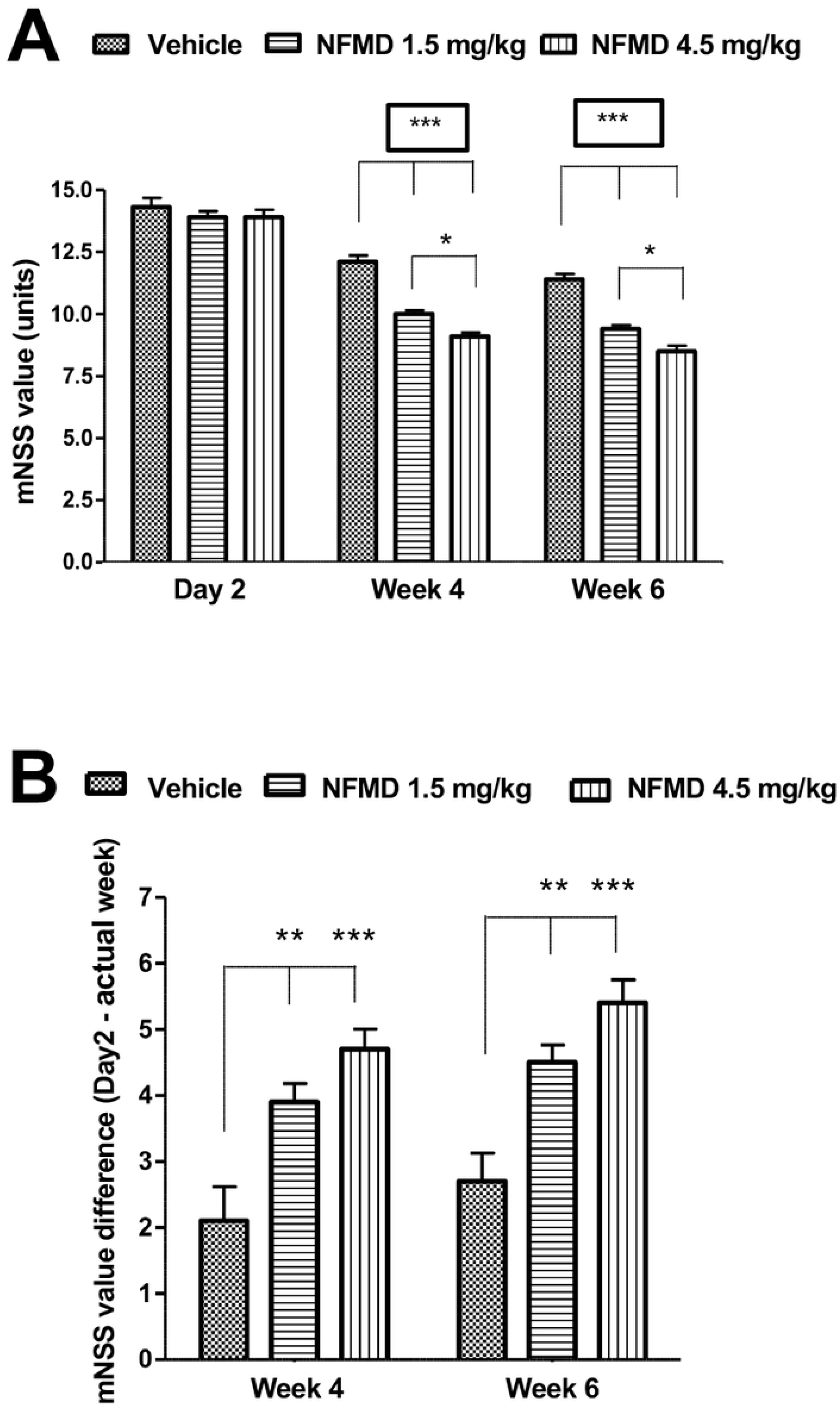
Dose-related neflamapimod (NFMD) subacute treatment effects on mNSS of rats with tMCAO. The study included three groups: vehicle (Pluronic F108; n=18), 1.5 mg/kg neflamapimod (n=21), and 4.5 mg/kg neflamapimod (n=21). (A) Mean±SD mNSS value at Day 2, Week 4 and Week 6. Both at Week 4 and Week 6, the mean±SD mNSS values were decreased and statistically significant lower in each neflamapimod dose group when compared to vehicle. In addition, a dose response effect on improvement in neurologic function was indicated by the lower mean±SD mNSS value in the 4.5 mg/kg dose group when compared to that of the 1.5 mg/kg dose group. (B) Absolute changes in mNSS±SD from Day 2 to Week 4 and Day 2 to Week 6. Statistically significant mNSS differences were observed at both time point comparisons for vehicle compared to 1.5 or 4.5 mg/kg dose groups. Significance of between group differences was assessed using a two-way analysis of variance (ANOVA) with a Bonferroni correction (\**P*<0.05; \*\**P*<0.01; \*\*\**P*<0.001).

Importantly, clear differences were demonstrated between the groups treated with neflamapimod compared to the vehicle-treated group. As shown in Fig 1A, the mean±SD mNSS values were statistically significantly lower (*P*<0.001 for all comparisons) in the 1.5 mg/kg dose group (10.0±0.7 at Week 4 and 9.4±0.7 at Week 6) and in the 4.5 mg/kg dose group (9.1±0.7 at Week 4 and 8.5±1.0 at Week 6) at both time points when compared to vehicle (see above, 12.1±1.1 at Week 4 and 11.4±0.9 at Week 6). These results document a positive effect of neflamapimod on neurologic repair following induction of ischemic stroke. Furthermore, a dose-related effect on improvement in neurologic function was indicated by the statistically significant lower mean mNSS in the 4.5 mg/kg dose group when compared to the 1.5 mg/kg dose group (*P*<0.05 at Week 4 and Week 6). Findings were similar when absolute changes in mNSS from Day 2 to Week 4 and Day 2 to Week 6 in the 1.5 mg/kg dose group (3.9 and 4.5, respectively) and the 4.5 mg/kg dose group (4.8 and 5.4, respectively) were compared to the absolute changes in the vehicle group (2.1 and 2.7, respectively), as summarized in Fig 1B. Statistically significant differences in absolute change in mNSS were observed at both time point comparisons for the 1.5 mg/kg dose group (*P*<0.01) and the 4.5 mg/kg dose group (*P*<0.001) compared to the vehicle control group.

It is also noteworthy that within all three individual groups the mean mNSS values were statistically significantly (*P*<0.001) decreased at the Week 4 and Week 6 time points versus Day 2. The vehicle group results for the mean±SD mNSS values (12.1±1.1 at Week 4 and 11.4±0.9 at Week 6 versus 14.3±1.7 on Day 2) imply a low degree of spontaneous recovery of neurologic functions in these study animals from Day 2 to Week 4 (*P*<0.001), as well as from Week 4 to Week 6 (*P*<0.05). It should be noted, however, that the mean mNSS value of the vehicle-treated control group decreased by ∼19% only during the six-week study period, and the mean mNSS score of 11.4 at Week 6 was nowhere close to the pre-stroke baseline mNSS value of 0 for healthy animals. Hence, the underlying spontaneous recovery process was very slow in this model.

### Neflamapimod-mediated improvement in motor and sensory function behavioral tests during the subacute stroke phase

Additional behavioral evaluations of motor and sensory functions (stepping test, body swing and forelimb placement) were performed to complement the mNSS end point for assessment of neflamapimod treatment effects during the subacute phase post stroke. The findings from these additional tests are presented in Fig 2 and were consistent with results of the mNSS evaluation.

**Fig 2.**
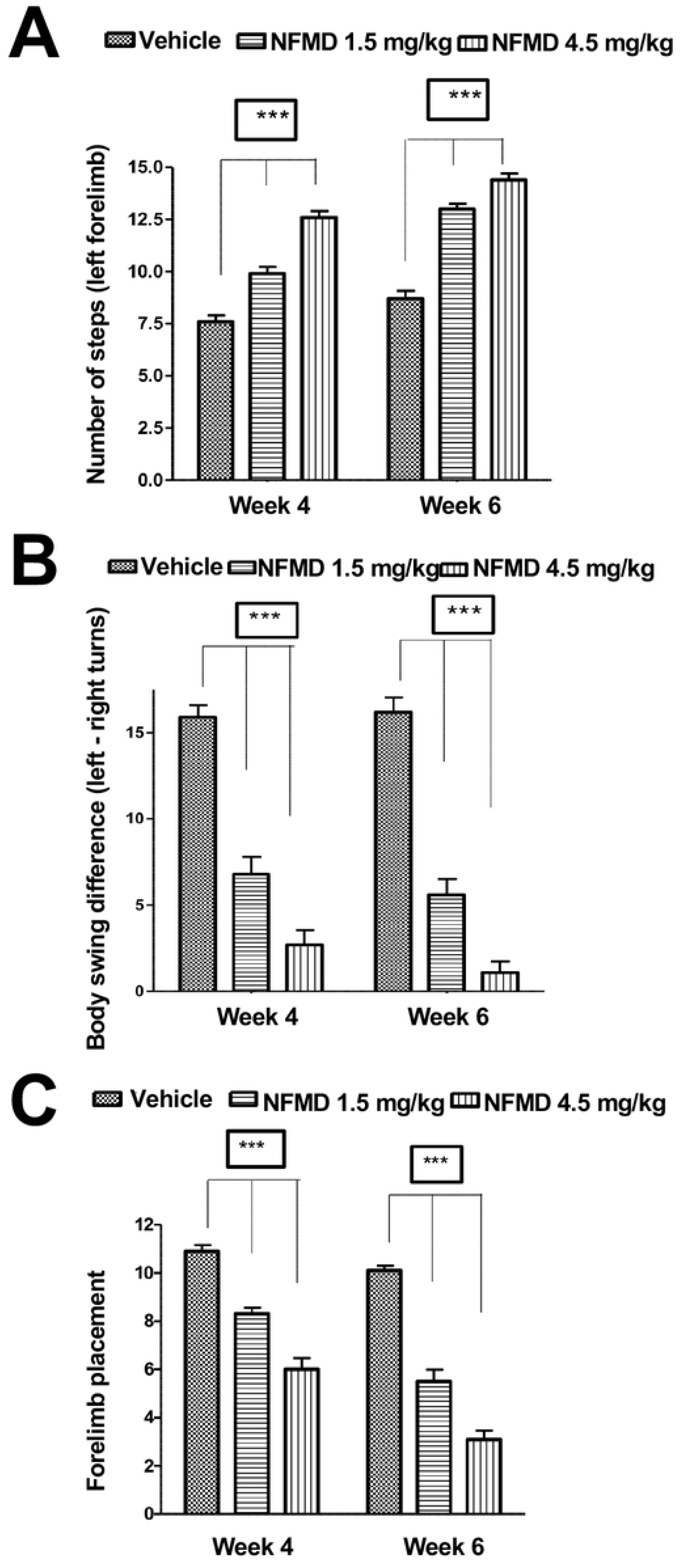
Effects of four- and six-week neflamapimod (NFMD) subacute treatment on stepping, body swing and forelimb placement tests of rats with tMCAO. Treatment groups include vehicle (n=18), 1.5 mg/kg neflamapimod (n=21), and 4.5 mg/kg neflamapimod (n=21). (A) Stepping test. Rats in both neflamapimod dose groups took a statistically significant greater mean±SD number of left forelimb steps at Week 4 and Week 6 compared to vehicle-treated animals. (B) Body swing test. Mean±SD adjusted body swing score values at Week 4 and Week 6 in the 1.5 mg/kg, 4.5 mg/kg dose groups were statistically significantly lower when compared to vehicle group. (C) Forelimb placement test. Mean±SD forelimb placement scores at Week 4 and Week 6 in the 1.5 mg/kg and 4.5 mg/kg dose groups were statistically significantly lower when compared to vehicle group. Statistical significance of between group differences was assessed using ANOVA with a Bonferroni correction (\**P*<0.05; \*\**P*<0.01; \*\*\**P*<0.001).

In the left forelimb stepping test, rats in both neflamapimod dose groups had a statistically significant (*P*<0.001 in all comparisons at a given time point) greater mean±SD number of steps at Week 4 (9.9±1.5 in 1.5 mg/kg group and 12.6±1.4 in 4.5 mg/kg group) and Week 6 (13.0±1.2 in 1.5 mg/kg group and 14.4±1.4 in 4.5 mg/kg group) compared to vehicle-treated animals (7.6±1.3 at Week 4 and 8.7±1.6 at Week 6) as shown in Fig 2A. Supporting that the effects of neflamapimod were specific to neurologic recovery following tMCAO, the mean±SD number of steps for the right forelimbs were similar in all groups at Week 4 (19.1±0.4 for 1.5 mg/kg, 19.1±0.4 for 4.5 mg/kg and 18.9.7±0.2 for vehicle group) and Week 6 (19.1±0.4 for 1.5 mg/kg, 19.2±0.4 for 4.5 mg/kg and 19.1±0.2 for vehicle group), respectively, and comparable to pre-stroke baseline left forelimb (19.8±0.6 for 1.5 mg/kg, 18.8±3.3 for 4.5 mg/kg and 19.7±0.8 for vehicle group) or baseline right forelimb (20.0±0.7 for 1.5 mg/kg, 20.1±0.7 for 4.5 mg/kg and 19.7±0.8 for vehicle group) values.

Additional results from the body swing (BSW) tests indicate a more prominent motor recovery in neflamapimod-treated animals. The mean±SD adjusted BSW score in the tests at Week 4 and Week 6 in the 1.5 mg/kg (6.8±4.6 and 5.6±4.2, respectively) and 4.5 mg/kg (2.8±3.9 and 1.1±2.9, respectively) dose groups were all statistically significantly (*P*<0.001) lower when compared to vehicle group (15.9±3.0 and 16.2±3.6, respectively) at the same time points, as presented in Fig 2B.

Finally, the mean±SD forelimb placement scores in the forelimb placement tests at Week 4 and Week 6 in the 1.5 mg/kg (8.3±1.2 and 5.5±2.2, respectively) and 4.5 mg/kg (6.0±2.1 and 3.1±1.7, respectively) neflamapimod dose groups were also statistically significantly (*P*<0.01 in all comparisons at the two time points) lower when compared to vehicle group (10.9±1.1 and 10.1±0.9, respectively), as shown in Fig 2C. The forelimb placement test scored the above-mentioned mean±SD forelimb placement scores positive values for the affected left forelimbs post stroke only. All the forelimb placement test scores for the left and right forelimbs at baseline, as well as all the values for the right forelimbs at Week 4 and Week 6 were zero. The lower forelimb placement scores observed in the neflamapimod-treated animals suggest a treatment effect of neflamapimod on recovery of somatosensory function following ischemic stroke.

### Study termination biomarker evaluation providing evidence for dose-dependent increase in brain BDNF protein levels mediated by neflamapimod treatment, and showing no significant effect on IL-1β production

To examine the potential effect of neflamapimod on relevant biochemical pathways following ischemic stroke, IL-1β and BDNF protein levels in homogenates of brain tissue harvested from the right (tMCAO side) and left (non-affected) hemisphere were determined by ELISA. Specifically, on Day 44, two days following the final dose of test article, brains were harvested from all animals and samples were processed after the brains were divided into left and right hemisphere. Samples from 18 animals in each treatment group (1.5 mg/kg neflamapimod, 4.5 mg/kg neflamapimod and vehicle control) were analyzed.

Interestingly, IL-1β levels were variably increased in a subset of animals in each group in the injured right brain hemisphere (Table 3). IL-1β was undetectable (i.e. below LLOQ; IL-1β=0 pg/ml) in the uninjured left brain hemisphere of all animals. The mean±standard error of the residual mean (SEM) IL-1β levels in the right hemisphere (after correction for an assay noise level of 5 pg/ml) were numerically lower in each neflamapimod dose group (14.6±5.1 pg/ml in the 1.5 mg/kg group and 15.2±6.7 pg/ml in the 4.5 mg/kg group) compared to the vehicle group (29.3±12.4 pg/ml), but there was no statistically significant difference between these groups. Thus, no clear neflamapimod effect on IL-1β could be detected on Day 44 post stroke.

**Table 3.** IL-1β (pg/ml) levels in the injured right brain hemisphere on Day 44 post stroke.

| Vehicle |  | 1.5 mg/kg NFMD |  | 4.5 mg/kg NFMD |  |
| --- | --- | --- | --- | --- | --- |
| Rat Number | IL-1β * (pg/ml) | Rat Number | IL-1β * (pg/ml) | Rat Number | IL-1β * (pg/ml) |
| 2 | 17.2 | 1 | 15.2 | 9 | 0 |
| 8 | 37.5 | 4 | 0 | 12 | 0 |
| 16 | 29.3 | 5 | 13.2 | 14 | 65.0 |
| 20 | 4.4 | 18 | 62.5 | 15 | 96.8 |
| 28 | 0 | 23 | 0 | 22 | 0 |
| 29 | 12.8 | 27 | 19.9 | 24 | 0 |
| 33 | 183.5 | 31 | 0 | 25 | 25.1 |
| 38 | 0 | 40 | 0 | 26 | 0 |
| 43 | 146.1 | 41 | 0 | 32 | 50.8 |
| 45 | 50.8 | 42 | 0 | 34 | 0 |
| 51 | 45.7 | 44 | 47.8 | 36 | 0 |
| 52 | 0 | 46 | 33.1 | 39 | 0 |
| 54 | 0 | 50 | 1.3 | 47 | 1.3 |
| 56 | 0 | 53 | 0 | 57 | 34.6 |
| 62 | 0 | 55 | 0 | 58 | 0 |
| 64 | 0 | 60 | 61.0 | 59 | 0 |
| 69 | 0 | 73 | 0 | 75 | 0 |
| 77 | 0 | 74 | 8.9 | 76 | 0 |
| **Mean** | **29.3** | **Mean** | **14.6** | **Mean** | **15.2** |
| **SD** | **52.6** | **SD** | **21.8** | **SD** | **28.5** |
| **SEM** | **12.4** | **SEM** | **5.1** | **SEM** | **6.7** |

\* IL-1β values after subtraction of the LLOQ value of 5 pg/ml.

Additionally, BDNF protein was detectable in both brain hemispheres of all animals on Day 44 post stroke as shown in Fig 3. Within each brain hemisphere there was a significant dose-related effect of neflamapimod for increasing BDNF protein levels (Jonckheere-Terpstra test: *P*<0.01 and *P*<0.05 for the left (Fig 3A) and right (Fig 3B) hemisphere, respectively). In addition, in the left hemisphere (non-affected side), the mean±SEM BDNF levels were statistically significantly higher in the 4.5 mg/kg group (2,762±271.6 pg/ml) than in the vehicle group (2,010±149.7; *P*<0.05). The left hemisphere mean±SEM BDNF levels in the 1.5 mg/kg group (2,213±136.0 pg/ml) were intermediate to those in the vehicle and the 4.5 mg/kg group, although not statistically significantly different from the vehicle group when the 1.5 mg/kg and vehicle groups were compared directly.

**Fig 3.**
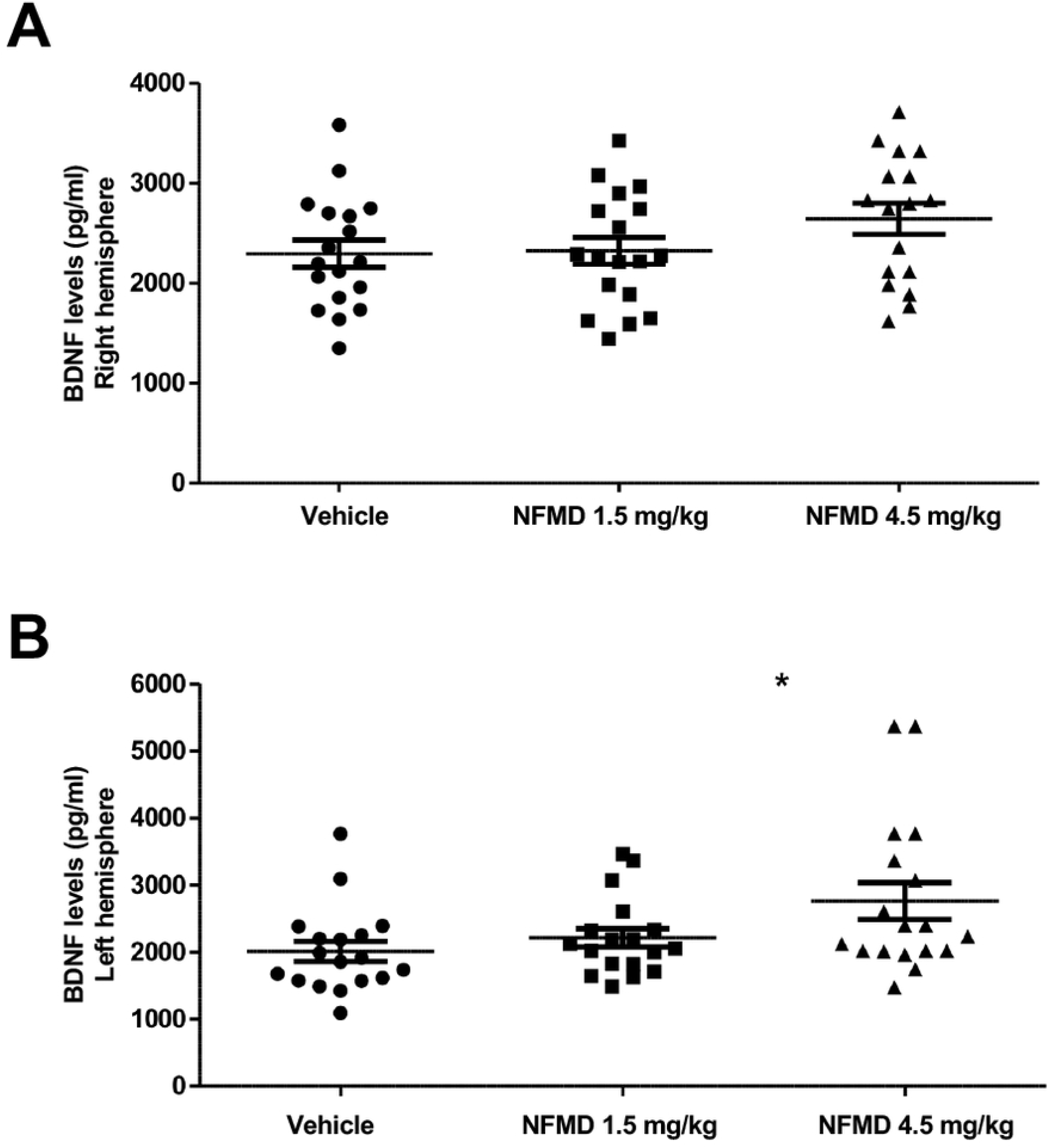
Dose-related neflamapimod (NFMD) subacute six-week treatment effects on rat brain BDNF levels analyzed on Day 44 post tMCAO. Brain homogenate samples from 18 animals in each treatment group (vehicle, 1.5 mg/kg neflamapimod, and 4.5 mg/kg neflamapimod) were prepared two days after the six-week treatment period and analyzed by BDNF ELISA. Data are presented as a technical data plot where the result for each individual animal is represented as one dot, the horizontal long line represents the mean of all values within a group, and the upper and lower range of the SEM are presented. (A) Right hemisphere (tMCAO): A significant neflamapimod dose trend by the Jonckheere-Terpstra test for higher BDNF levels was observed (*P*<0.05) for across the three groups. (B) Left hemisphere (non-injured side): The Jonckheere-Terpstra test indicated a significant neflamapimod dose trend for higher BDNF levels in the left hemisphere (*P*<0.01). The mean±SEM BDNF level was also statistically significantly higher in the 4.5 mg/kg neflamapimod group when compared with the vehicle control group (*P*<0.05).

## Discussion

The results for the daily general health assessments, the weekly body weight measurements, the neurologic recovery assessments on Day 2, at Week 4 and at Week 6 post stroke, and the terminal biomarker measurements on Day 44 in this rat tMCAO study taken together led to the following seven key observations for evaluation of delayed and prolonged neflamapimod treatment. (1) Neflamapimod treatment dose-dependently and continually improved neurologic and behavioral outcomes assessed by mNSS and specific measures of sensory and motor function post stroke (Fig 1 and Fig 2). (2) Neflamapimod dose-dependently increased terminal brain BDNF protein as a neurogenic factor biomarker measure for beneficial effects on synaptic plasticity (Fig 3). (3) Neflamapimod and vehicle treatments were generally well-tolerated throughout the study, providing no treatment-related adverse clinical signs. (4) There was an average body weight gain of 27% in all treatment groups throughout the six-week study period (Table 2) and there was no difference observed in the general health of the rats that would have contributed to any neflamapimod versus vehicle treatment-mediated functional recovery. (5) Slight, albeit statistically significant spontaneous recovery over time was also observed in the vehicle control group (an average ∼19% decrease in mNSS value within six weeks; Fig 1) implying that neflamapimod *in vivo* efficacy may enhance a biologic process, like neural or synaptic plasticity, that is already active. (6) Generally, the functional recovery was slow and steady in all neflamapimod- or vehicle-treated animals throughout the prolonged study period (Fig 1 and Fig 2). (7) There were signs for unresolved IL-1β-mediated chronic inflammation in the injured brain hemisphere of a subset of animals, however, no clear neflamapimod effect on IL-1β production was apparent at study termination (Table 3). Taken together, the results for this research study provide evidence that there is an additional clinical opportunity for p38α inhibitors in ischemic stroke since neflamapimod widened the window to treatment initiation to 48 hours post stroke in a rat tMCAO model and promoted recovery.

The most important finding of this study is that the delayed and prolonged treatment with neflamapimod resulted in dose-related neurologic and behavioral improvements versus vehicle control, when this p38α inhibitor was administered orally to rats with tMCAO in an experimental paradigm of promoting synaptic plasticity and functional recovery during the subacute phase of ischemic stroke. To this end, neflamapimod treatment started at 48 hours after reperfusion at a time when the acute stroke phase was considered complete, since this represents a time point that is outside the previously characterized neuroprotection window of less than 24-hours post tMCAO-induced stroke for a p38α inhibitor [25]. In the most relevant (based experimental stroke model and rat strain) previously published study, the p38 MAPK inhibitor SB203580 was administered via intracerebroventricular injection 30 minutes pre-tMCAO, or at 6, 12 or 24 hours post-tMCAO to Sprague Dawley rats and demonstrated a decrease in infarct volume that was associated with anti-inflammatory effects in the injured brain and improved recovery from neurologic deficit measured on Day 2 post stroke for all the treatment initiations up to 12 hours post-tMCAO, whereas no beneficial effect was observed for the treatment initiation at 24 hours post tMCAO [25]. The observation that a 12-hour time window for the start of p38α inhibition, but not a 24-hour window allowed for neuroprotective effects in the rat tMCAO-induced stroke model supports the selection of 48-hours as the time point for initiation of subacute phase neflamapimod dosing in the current study, since the results by Piao *et al*. [25] indicated that it is too late a time point for a p38α inhibitor in this type of model to lead to reduction of neuronal loss as a mechanism for impacting functional outcome.

This neflamapimod treatment study is clearly different from all previously reported studies of other small molecule p38 MAPK inhibitors in various experimental stroke models [25, 28-32] both in terms of the delay in treatment initiation to 48 hours post reperfusion and the six-week prolonged dosing duration and overall treatment study focus. All previously reported studies of p38 MAPK inhibitors that most potently inhibit the p38α isoform, including SB203580 [25], SB239063 [28, 29], RWJ67657 [31], or VCP979 [32], involved compound administration before or soon after the experimental stroke during the acute phase, with treatment initiations at different time points, ranging from as early as 1 hour pre stroke to as late as 24 hours post stroke. Additionally, all prior p38α inhibitor studies had a focus on demonstrating neuroprotective effects that resulted in significant reduction of infarct volume (assessed as early as 6 hours post stroke) and subsequent improvements of neurological defects assessed at 24 hours [28, 29], 2 days [25], 7 days [32] or 14 days post stroke [31]. The two most recent p38α inhibitor studies [31, 32] were also the two longest ones prior to the present study. Interestingly, they did demonstrate the validity of mNSS [51] as a readout for functional recovery attributable to neuroprotection via p38α inhibition, that was measurable during the subacute phase after photothrombotic ischemic stroke in mice at one and two weeks, respectively [31, 32]. The results of this neflamapimod study extend these positive mNSS effect observations for a p38α inhibitor to a rat tMCAO model. In contrast to all previous p38α inhibitor studies, the actual size of the acute cerebral infarct was not measured in this neflamapimod study since the main objective was to evaluate the compound as a therapy for post stroke functional recovery, not as a neuroprotection therapy, and the delay in treatment initiation to 48 hours post stroke was expected to prevent a neuroprotection immediate impact of the compound on infarct size based on the previously published p38α inhibitor results [25]. Therefore, infarct volume measurement was not relevant for evaluation of neflamapimod due its late application as experimental stroke treatment. Instead, evaluation of mNSS [51] and additional functional tests (stepping test, body swing test, forelimb placement test) to assess the sensory motor deficit after stroke together with BDNF protein biomarker effects were deemed sufficient to evaluate and demonstrate the beneficial *in vivo* efficacy of neflamapimod.

Since the present study represents a first both for initiating the dosing of a potent and selective p38α inhibitor as late as 48 hours in the subacute phase after rodent tMCAO-induced stroke, as well as for dosing this type of a small molecule for a prolonged time period of six weeks, it is a challenge to find directly supportive subacute or chronic stroke phase studies. It is reassuring that the functional recovery demonstrated in this study is consistent with a report by Umezawa *et al*. [52] who demonstrated that selective inhibition of p38α by SB239063 improved locomotor recovery after SCI in mice. Although in a different disease context, this study may be most relevant since Umezawa *et al*. [52] also utilized a genetic approach (i.e. a heterozygous knockout of MAPK14) to identify the p38α specificity of the effect. Generally, small molecule treatments in the rodent cerebral ischemia model start before the 48-hour time point after reperfusion, as already discussed for all the previously reported p38α inhibitor studies [25, 28-32]. Those studies are not inconsistent with this neflamapimod study, only different in that the neurologic recovery effects therein are clearly associated with a primary acute phase neuroprotective effect of the p38α inhibitors [25, 28-32]. A review of the experimental stroke literature, however, revealed that the results of all these studies are in conflict with several reports that have associated elevated p38 MAPK activity, including p38α activity, with a neuroprotective role during and after stroke, either in the acute or subacute phase [53-59]. One caveat is that several of these differing studies used the small molecule SB203580 at a relatively high concentration and did not address the possibility of blocking p38 MAPK isoforms other than p38α, such as p38β, or other kinases (e.g. GAK, RIPK2, NLK, JNK3, CSNK1D and others) with this less potent and much less selective p38α inhibitor tool compound [60]. As an example, SB203850 through inhibiting casein kinase Iδ and ? blocks WNT-stimulated β- catenin signaling [61, 62], which might account for compound effects on neurogenesis, therefore, render claims for p38 attributed activities invalid [56]. While it is likely that the role of p38α depends on context, which may include the model species or strain, the type of stroke model, the temporal characteristics, the local concentration of the target protein and its substrates, the target cell type, the presence or absence of pathway regulators, and possibly other factors, it appears that additional experimental analyses involving low concentrations of neflamapimod would have been useful to clarify the role of p38α in some of these conflicting research studies. Based on its low selectivity entropy score in addition to suitable physical, biochemical and cellular properties, neflamapimod has been recommended as the small molecule compound to utilize in experimental studies that have the objective of understanding the biologic effects of inhibiting p38α kinase activity specifically [63].

The rat tMCAO model used in this study is one of the most frequently used *in vivo* models in stroke research, partially because the procedure used for stroke induction causes reproducible infarcts in the middle cerebral artery [33, 64]. The tMCAO procedure was performed under highly standardized and well-controlled conditions [65] by well-trained and experienced scientists to reduce the inter-animal variability of infarct size to a minimum, which is demonstrated by consistently high mNSS values determined on Day 2. The mean±SD mNSS value was 14.0±1.4 for the 67 rats prior to their randomized assignment into the three different treatment groups. This a relatively tight value and an indication of a robust stroke since it represents a tremendous increase from the mNSS value of 0 reflecting the normal, healthy pre-tMCAO status prior to Day 1.

Even though the technical procedures used for this experimental model have been designed to have a low mortality associated with the robust stroke induction, mortality of up to 25% within 24 hours after stroke onset is quite common [66-68] and mortality has been reported to increase up to 33% in the following days due to edema [69, 70]. In this tMCAO study, 9 out of 76 animals (∼12%) were lost during the first 24 hours post stroke, 4 more until 48 hours post stroke, and 3 more during the first two weeks of treatment, resulting in ∼21% mortality within the first 16 days after tMCAO. These results are well within the range of the expected mortality results for rodent stroke models [66-70]. Intriguingly, the mortality results in these experimental models also somewhat mirror human data where mortality within 28 days of stroke has been described to range up to ∼28% [71, 72].

Oral neflamapimod was dosed at 1.5 and 4.5 mg/kg administered in 1% Pluronic F108 vehicle twice daily on 5.5 days per week for six weeks in the present rat tMCAO study. These two neflamapimod dose levels in this vehicle were selected because their twice daily administration for three weeks was previously demonstrated to be pharmacologically active in aged rats with cognitive deficits attributed to chronic inflammation-induced, IL-1β-mediated impairment of synaptic plasticity [47]. The results obtained in this tMCAO study corroborate that these two neflamapimod dose levels are pharmacologically active in rats. Unlike the aged rat study, in which the 4.5 mg/kg dose did not lead to better cognitive results compared to the 1.5 mg/kg dose [47], the beneficial effects by neflamapimod resulting in neurologic and behavioral improvements after stroke increased in a dose-related manner and enhanced the slight spontaneous recovery observed in the vehicle control group. Similar to the aged rat study [47], the beneficial effects of neflamapimod in this experimental stroke study do not appear to be due to inhibition of IL-1β production. There was no statistically significant decrease in the terminal IL-1β brain levels at either neflamapimod dose level compared to vehicle control.

The IL-1β results of the present rat tMCAO study are limited to one time point only and show that a residual elevation of IL-1β on the stroked side of the brain, but not on the unaffected side, was measurable in 50% of animals in the vehicle- and low-dose neflamapimod-treated group as well as in 30% of the animals in the high-dose neflamapimod-treated group at the end of the study. This together with the observed beneficial neurologic effects by neflamapimod suggests that chronic inflammation may play a role in this rat tMCAO stroke model. This is a finding that supplements previously reported IL-1β results in this type of model [36, 73, 74]. Although this single time point analysis appears to exclude a neflamapimod effect on IL-1β production, it does not necessarily preclude a neflamapimod effect on IL-1β signaling, even though a correlative relationship of the Day 44 IL-1β levels with the six-week mNSS values or with their Day 44 BDNF levels was not found. The present study was not intended to address the exact mechanism of action of neflamapimod and the IL-1β and BDNF data are very limited.

IL-1β is a pleiotropic cytokine that is known to be upregulated acutely after ischemic stroke in the rat and has been demonstrated to be a contributor to pathology via p38α activity [25]. Previously published temporal profiles of IL-1β mRNA or protein in experimental stroke, which are generally limited to a time period up to 5 days post stroke, have demonstrated that IL-1β is increasing shortly after stroke, usually peaking within the first 24 hours during the acute phase, and then declining towards baseline levels or still remaining elevated [36, 73, 74]. While it is clear that acutely elevated levels of IL-1β exert potent pro-inflammatory functions and play a role in initiation of tMCAO-induced damage [75, 76], less is known about IL-1β dynamics as a result of stroke damage, or about what goes on mechanistically during the subacute stroke phase [37]. The results of the present study are in line with studies that demonstrate that subacute phase IL-1β continues to potentiate neuronal injury in experimental stroke [37]. They run counter to other studies that argue that subacute/chronic IL-1β after experimental stroke serves to halt or repair injury [37]. It is possible that these controversial results may be context-dependent (e.g. species/strain, age of animals, experimental stroke model, size of stroke damage, involved cell types) and additional research will be needed to sort out the relevance of the different outcomes and the mechanisms for IL-1β-mediated responses [37]. It is clear that modeling inflammation and immunity in experimental models of stroke is challenging even though the sensitivity of rodents to focal cerebral ischemia as well as their physiology and pathophysiology are sufficiently similar to humans to make them a highly relevant model organism [77, 78].

Terminal measurements of brain BDNF protein demonstrated that six-week neflamapimod treatment resulted in dose-dependent increases of this neurotrophic factor in both the injured and uninjured hemisphere. BDNF is a key regulator of plasticity both in the healthy and injured brain which has been recognized as a key regulator of rehabilitation- and activity-induced functional and motor recovery, respectively, after stroke [79-81]. BDNF is reported to have a critical role in promoting recovery after stroke as a crucial signaling molecule that mediates adaptive brain plasticity [35, 82-85]. Increased BDNF levels in perilesional areas have been observed with interventions that improve functional recovery post stroke [86-88]. Conversely, attenuation of brain BDNF levels or effects following cerebral ischemia results in reduced neuroplastic changes or decreased recovery of function, both in spontaneous and in rehabilitation-induced recovery scenarios [84, 89, 90]. Since regenerative roles have been attributed to BDNF in preclinical models of stroke [84, 91-94], upregulation of BDNF may be a plausible contributor to the neflamapimod-induced functional recovery observed after the ischemic stroke.

How BDNF elevation is linked to p38α inhibition by neflamapimod in this experimental stroke model will need to be determined in a follow-up mechanistic study. Our favored hypothesis is that the BDNF levels might be interpreted as a marker of a more general effect on IL-1β signaling [42] that could result from p38α inhibition [41, 43]. In such a hypothetical mechanistic model, the observation of inhibition of IL-1β signaling without impacting IL-1β production would be consistent with previously observed aged rat results demonstrating that neflamapimod is more potent at inhibiting IL-1β signaling than at inhibiting IL-1β production [47]. While this stroke study was not intended to address the exact mechanism of action of neflamapimod, the observation that BDNF protein was increased in both brain hemispheres at study termination is nevertheless supporting the underlying hypothesis that functional recovery was associated with enhancing synaptic plasticity.

Taking into account the design of the study that precludes an effect through neuroprotection, the absence of an anti-inflammatory effect, and the possibility of an effect on IL-1β signaling, combined with the scientific literature regarding p38 MAPK-mediated deleterious effects of IL-1β on synaptic plasticity [43, 95], the results herein imply a model in which IL-1β would limit functional recovery after stroke via p38α-mediated impairment of neural and synaptic plasticity. The results in this neflamapimod-mediated stroke recovery study are also consistent with studies in other disease contexts in which activation of p38 MAPK, particularly the alpha isoform, is associated with impaired synaptic plasticity [18, 96]. Further, the non-clinical to clinical translational potential with neflamapimod as a p38α inhibitor is that, similar to the observed slight spontaneous recovery observed in the vehicle control group, neural and synaptic plasticity has been argued to be active in humans and at least partially effective in recovery after stroke [97]. Enhancing a biologic process like neural or synaptic plasticity, that is already active, rather than targeting a process that may not represent an intrinsic recovery pathway, such as neurogenesis, should at least theoretically be more likely to have a clinical effect. For this reason, the opportunity of translational success with p38α inhibition to promote recovery following stroke by enhancing plasticity would be expected to be higher than for neuroprotection.

This research study can also be considered as a non-clinical development study that assesses the *in vivo* efficacy of neflamapimod, a highly selective p38α inhibitor, in the commonly used rat tMCAO model of stroke [33, 64]. Even though the promptly restored blood flow in this experimental model is different from the pathophysiology in spontaneous human stroke, the rat tMCAO model closely mimics the therapeutic situation of mechanical thrombectomy which is expected to be increasingly applied to patients [33]. This suggests that this rat ischemia model may have clinical relevance [33].

Potentially the most important consideration for clinical translation when targeting recovery from stroke is the practical consideration of time window for therapeutic intervention. The narrow time window after onset of ischemia that is required of neuroprotective approaches has posed an insurmountable challenge for clinical development [98]. Stroke patients do not often present within the required first few hours and, even within the initial 24 hours post-stroke, the clinical presentation does not allow one to assess the true size and severity of the stroke. Therefore, any clinical study that starts treatment of patients within the first day of the stroke has a highly variable and heterogeneous patient population and thus requires a large sample size to demonstrate clinical effects. These restrictions preclude the ability to demonstrate clinical proof-of-concept in phase 2 clinical testing. In contrast, a time window of 24 to 48 hours after stroke allows patients’ clinical course to have stabilized and allows time for a clinical exam and a diffusion-MRI scan to precisely determine the location and extent of the stroke before starting treatment. In addition, a reasonable prognostic indication of the extent of recovery that a patient will attain with or without intervention can be assessed. As a result, starting treatment in a clinical study at 48 hours or later after stroke allows for the inclusion of more homogenous sub-populations of patients and increases statistical power with fewer subjects; therefore, definitive clinical-proof-of-concept could potentially be demonstrated within a phase 2 clinical study.

Despite the observation of robust *in vivo* efficacy of neflamapimod, there are some limitations to this research stroke study, since only one treatment duration was chosen, and functional outcomes were measured at the Week 4 and Week 6 timepoints after the stroke. It cannot be predicted whether longer neflamapimod treatment duration with additional analyses would have led to further mNSS improvement and continued recovery during the chronic phase, and the exact timing of the onset of neflamapimod action on functional recovery was not determined. Neflamapimod treatment was initiated at 48 hours after reperfusion, at a time point that was considered subacute post stroke in light of the prior negative experiences with other p38α inhibitors and numerous neuroprotective agents when they were administered at 24 hours or more post-stroke, which implies an exceedingly low likelihood that neuroprotection plays a role in promoting functional recovery, but without accompanying histologic analyses an effect of neflamapimod on neuronal loss cannot be formally excluded. Further, while we believe that it is highly probable that the functional recovery mediated by neflamapimod was due to enhancement of plasticity mechanisms, in the absence of technologies that can measure synaptic function *in vivo* in real-time, there is no means to directly verify that assumption. Similarly, while the known activity of IL-1β to impair synaptic plasticity via p38α implies that inhibition IL-1β activity is a major contributor to the neflamapimod clinical activity, there is no means to directly confirm that link *in vivo*, or to exclude other potential mechanisms. Finally, in the current study, young (3-months old) male animals were utilized. Although neural and synaptic plasticity recovery functions appear to be active in aged animals and are also at least partially preserved in elderly patients [99], in order to improve clinical translation, a replication and extension of this study to include females, aged animal and animals with co-morbidities may be required since gender, age and reduced health condition (e.g. illnesses, diseases, disorders, health problems) may affect development of ischemic damage and resulting behavioral deficits in patients [100].

The clinical science translational opportunity with neflamapimod is that the compound has already been studied in early Alzheimer’s disease patients at the human dose equivalents of 1.5 and 4.5 mg/kg in the rat [46]. These doses were well-tolerated, achieved target cerebral spinal fluid (CSF) drug levels consistent with robust blood-brain-barrier penetration, and demonstrated target activity as assessed by imaging and CSF biomarkers. Thus, neflamapimod provides the opportunity of evaluation in the clinic to address whether its p38α inhibition will promote recovery after ischemic stroke, and such a study expected to provide insights into its potential to enhance neural and synaptic plasticity mechanisms in humans.

## Conclusions

Here prolonged, six-week oral administration of the p38α inhibitor neflamapimod, with treatment initiation starting at 48 hours post reperfusion that is outside the previously characterized neuroprotection window for p38α inhibitors, resulted in dose-related significant neurologic recovery and improvement of motor and sensory functions measured at four and six weeks post stroke. Additionally, dose-related increases of the neurogenic factor BDNF in the brain as a potential biomarker for neflamapimod effects on synaptic plasticity were observed at termination of the study on Day 44. Thus, neflamapimod use offers the possibility of widening the window for treatment initiation post stroke and promoting recovery after a completed stroke. Since it is also known to counteract cognitive decline in aged rats a well as Alzheimer patients, neflamapimod is clinically developable and especially appealing as potential stroke therapy.

## Acknowledgments

The authors would like to acknowledge the help of all the people involved in this project, in particular Dr. Itschak Lamensdorf at Pharmaseed, who provided inspiration and interpretive perspective on the results. Additionally, Molly Opferman as consulting medical writer provided technical writing assistance in preparing the manuscript.

